## SupplementalFile for "Deterministic Early Endosomal Maturations Emerge From a Stochastic Trigger-and-Convert Mechanism"

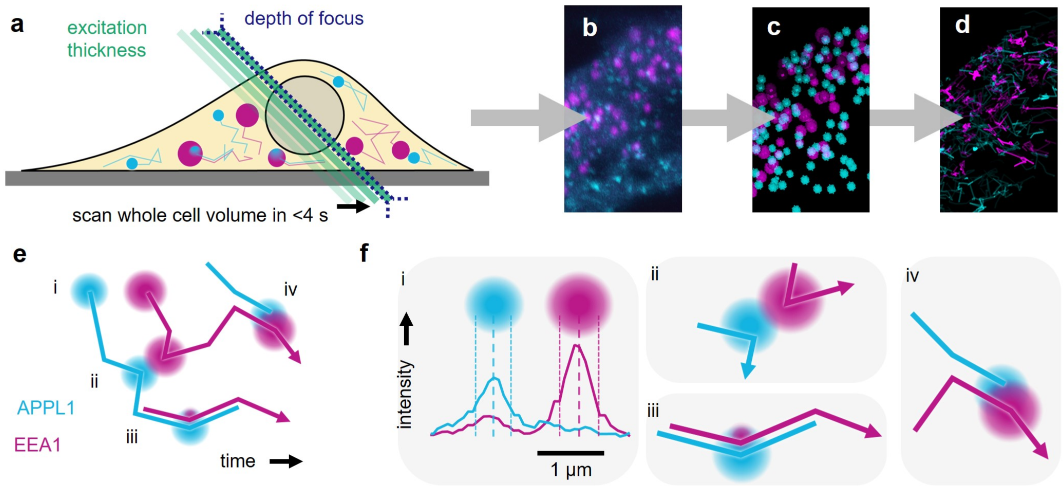


**Fig. 1. Image analysis pipeline to measure ensemble endosome characteristics. (a)** APPL1-EGFP and TagRFP-T EEA1 were imaged using LLSM at 2.5 s/volume. **(b)** Representative image of raw data, representing a single zoomed frame of a whole cell volume lasting up to ∼30 minutes. **(c)** Preliminary endosomes in each channel were identified separately by blob detection then an unsupervised pattern recognition-based routine was used to identify true endosomes, following by **(d)** linking based on localisation and intensity values to construct complete trajectories with continuous spatial data and intensity traces. **(e)** Trajectories of endosomes identified from opposite channels were then analysed together to identify of events of interest (i–iv). **(f)** Schematic showing the protocols used to detect each type of event. (i) The intensity profile between two endosomes (*cyan*: APPL1, *magenta*: EEA1) along the line connecting their centres of mass is shown. Dashed line indicates centre of mass; dotted lines indicate endosome boundaries determined by detection routine. Changes in surface-to-surface distances between pairs of nearby events are used to identify events, with (ii) collisions defined as the point of nearest approach, if below the threshold for surface-to-surface separation and (iii) conversions calculated by identifying regions of colocalised trajectories; (iv) such events are considered to be fusions if the EEA1 track that remains after the disappearance of the APPL1 track existed as a distinct tracked object prior to the colocalisation event.

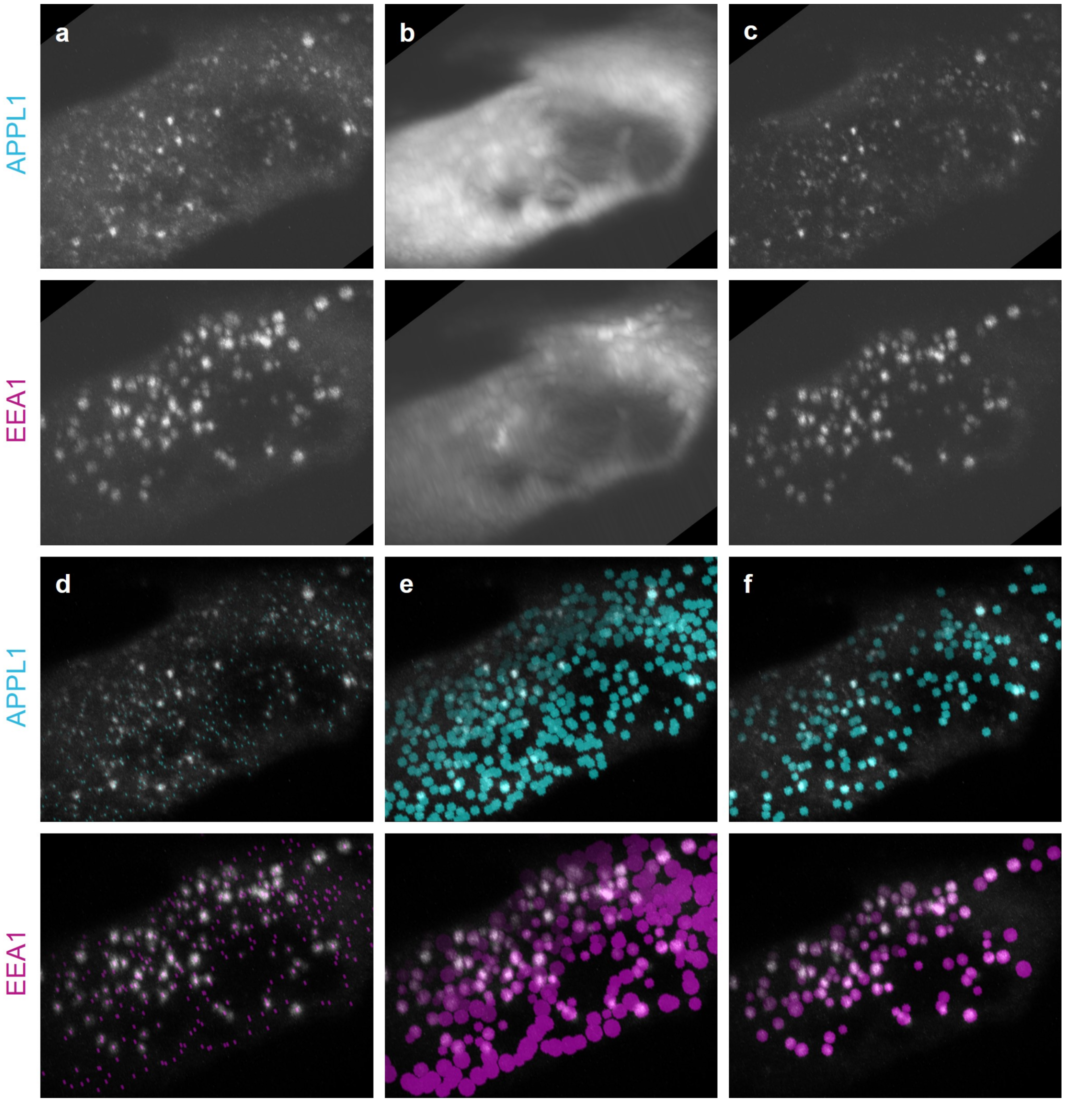


**Fig. 2. Schematic of full image analysis routine.** Overview of steps in image analysis routine. Each panel shows APPL1 (*top*) and EEA1 (*bottom*) channels, which are analysed independently of each other. **(a)** Deskewed images. **(b)** Local background is calculated by removing large bright objects from the raw data, then calculating the mean-intensity projection over time for all unmasked pixels. **(c)** Subtraction of local background improves signal-to-noise and is used as an input to a blob detection algorithm, which returns **(d)** centres of mass and **(e)** approximate radii for all potential endosomes. The initial threshold is set so as to favour over-detection, while also prioritising separation of nearby endosomes into distinct objects. **(f)** Final set of labelled endosomes. Blobs are clustered into two populations using a $k$-means clustering algorithm with an automatically calculated set of features that describe true endosomes and background, respectively, for each individual movie. Visual inspection confirmed that this approach captures both obvious (large and bright) as well as non-obvious (small and/or dim) endosomes, while separating nearby endosomes into distinct objects (which is important for accurate calculation of collisions and fusions).

| Identification of cotracking events | |
|---|---|
| Minimum number of contiguous cotracking frames | 4 frames |
| Maximum surface-to-surface distance | 0 µm |

| Identification of conversion events | |
|---|---|
| APPL1 track started before cotracking event | $\geq$ 30 s |
| EEA1 track continued after cotracking event | $\geq$ 1 frame |
| APPL1 track stopped at end of cotracking event | same frame |
| Average APPL1 intensity change from first to last 20% of event | decrease |
| Average EEA1 intensity change from first to last 20% of event | increase |

| Identification of collision events | |
|---|---|
| Relative local minimum in surface-to-surface distance | 25% |
| Absolute local minimum in surface-to-surface distance | $<$200 nm |
| Absolute separation between local minima in surface-to-surface distance | 3 frames |
| Absolute global maximum in surface-to-surface distance | $\geq$500 nm |

| Identification of fusion events | |
|---|---|
| EEA1 track started before cotracking event | $\geq$ 1 frame |

**Table 1. Parameters to automatically label events.** Cotracking events were defined as at least 4 contiguous frames from independently identified APPL1 and EEA1 tracks, where the surface-to-surface distance did not exceed 0 µm. Conversion events suitable for time series analysis required APPL1 signal to be detected for at least 30 s prior to the initiation of cotracking, EEA1 signal to be detected at least 1 frame after cotracking, and APPL1 signal to not be detected after cotracking. Successful conversions were considered those where APPL1 (EEA1) intensity decreased (increased), on average, between the first and last 20% of each event. Heterotypic collision events were first identified by computing the surface-to-surface distance between all neighbouring APPL1 and EEA1 tracks, with collisions identified as local minima in the time course of the surface-to-surface distance with both relative and absolute thresholds of 25% and 200 nm, respectively, and having a global maximum distance that at some point during tracking exceed$>$500 nm. In pairs of APPL1 and EEA1 tracks with multiple potential collisions, at least 3 frames of separation were required between local minima. Finally, cotracking events were segmented into fusions and conversions based on prior existence of EEA1 track: if the EEA1 track existed prior to APPL1–EEA1 cotracking, the event was classified as a fusion.

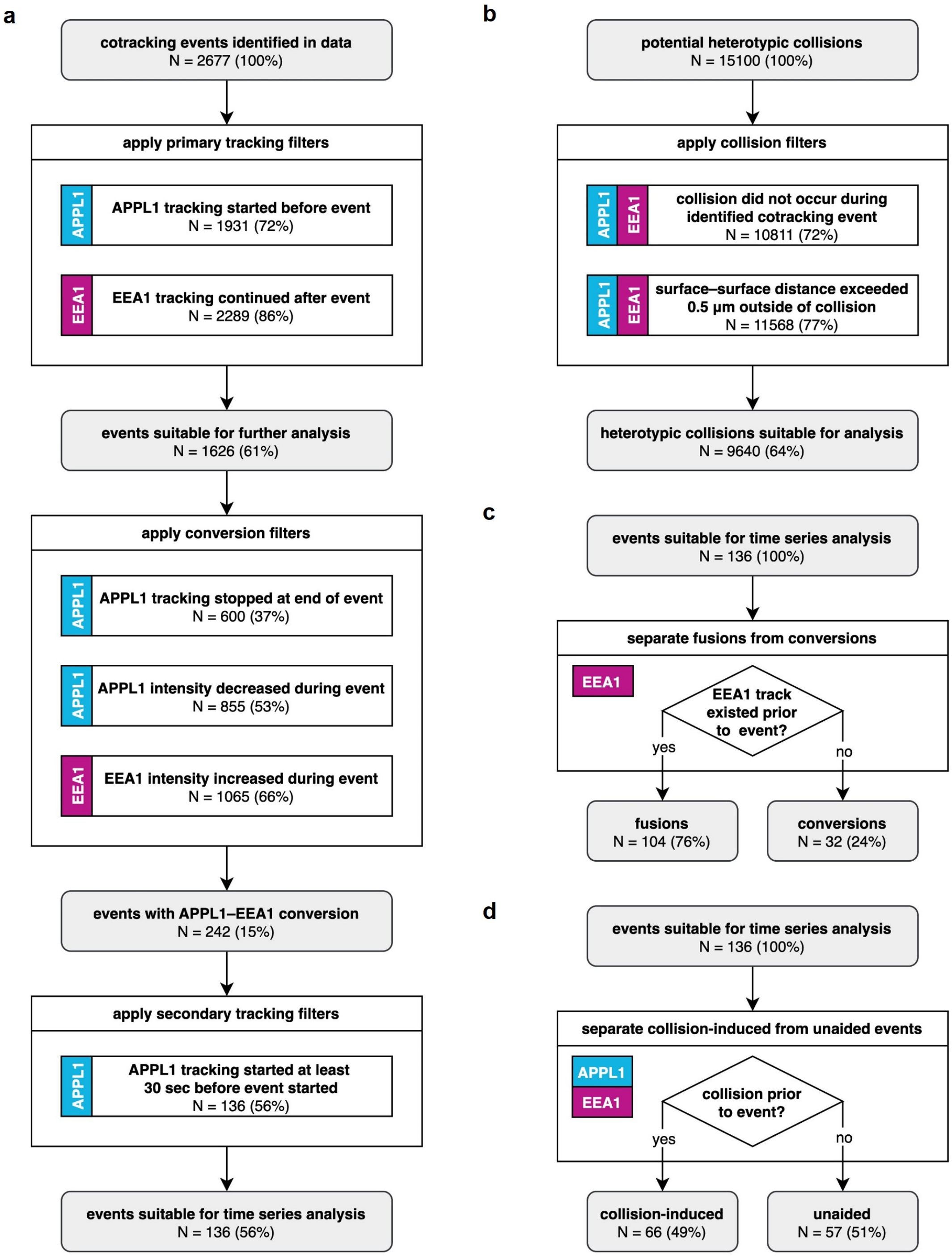
a
cotracking events identified in data
N = 2677 (100%)
apply primary tracking filters
APPL1
APPL1 tracking started before event
N = 1931 (72%)
EEA1
EEA1 tracking continued after event
N = 2289 (86%)
events suitable for further analysis
N = 1626 (61%)
apply conversion filters
APPL1
APPL1 tracking stopped at end of event
N = 600 (37%)
APPL1
APPL1 intensity decreased during event
N = 855 (53%)
EEA1
EEA1 intensity increased during event
N = 1065 (66%)
events with APPL1–EEA1 conversion
N = 242 (15%)
apply secondary tracking filters
APPL1
APPL1 tracking started at least
30 sec before event started
N = 136 (56%)
events suitable for time series analysis
N = 136 (56%)
b
potential heterotypic collisions
N = 15100 (100%)
apply collision filters
APPL1
EEA1
collision did not occur during
identified cotracking event
N = 10811 (72%)
APPL1
EEA1
surface–surface distance exceeded
0.5 µm outside of collision
N = 11568 (77%)
heterotypic collisions suitable for analysis
N = 9640 (64%)
c
events suitable for time series analysis
N = 136 (100%)
separate fusions from conversions
EEA1
EEA1 track
existed prior
to event?
yes
no
fusions
N = 104 (76%)
conversions
N = 32 (24%)
d
events suitable for time series analysis
N = 136 (100%)
separate collision-induced from unaided events
APPL1
EEA1
collision prior
to event?
yes
no
collision-induced
N = 66 (49%)
unaided
N = 57 (51%)

**Fig. 3** ***(previous page)*****. Workflow to automatically label events.** A series of filters were applied to the results of the trajectory analysis to select only events meeting stringent criteria. **(a)** Conversion events were identified by several increasingly stringent filter steps. Firstly, all cotracking events were found, taking as inputs the centres of mass and radii calculated in the image analysis routine. Cotracking events were defined as at least 4 contiguous frames from independently identified APPL1 and EEA1 tracks, where the surface-to-surface distance was $<0$ µm. Primary tracking filtering steps were firstly applied based on meeting 'yes'/'no' criteria, retaining only cotracking events having independent APPL1 trajectories that started before, and EEA1 trajectories that continued after, each event. Next, both tracking- and intensity-based filtering steps were applied to select successful conversions from all cotracking events. Conversions were defined as events where (i) APPL1 tracking stopped at the end of APPL1–EEA1 cotracking, (ii) APPL1 intensity decreased, on average, between the first and last 20% of each event, and (iii) EEA1 intensity, on average, between the first and last 20% of each event. By this stage, 9% of cotracking events were identified as conversion events. Finally, secondary tracking filtering steps were applied to ensure sufficient data points for time series analysis, retaining only trajectories for which APPL1 signal was detected for at least 30 s prior to the initiation of APPL1–EEA1 cotracking. 5% of colocalisations qualified as examples of completed APPL1–EEA1 conversion events appropriate for time series analysis. **(b)** Potential heterotypic collision events were first identified by computing the surface-to-surface distance between nearest neighbours of APPL1 and EEA1 over time, and finding the local minima in this time course where (i) the absolute distance was $<200$ nm of separation and (ii) the separation from the nearest local minimum in the time course of the surface-to-surface distance was at least 3 frames. These data were further filtered by two methods to separate potential cotracking events, rejecting all events that were (i) previously identified as part of cotracking events in **a**, and (ii) not having a surface-to-surface distance that at some point during tracking exceed 500 nm of separation. 64% of potential heterotypic collisions qualified as appropriate for consideration for events analysis. **(c)** Cotracking events were segmented into fusions and conversions based on prior existence of EEA1 track. If the same EEA1 endosome was independently tracked prior to APPL1–EEA1 conversion, the event was classified as a fusion. **(d)** Cotracking events were segmented into collision-induced and unaided events based on existence of any APPL1–EEA1 collision in the 30 s before the event. Events (conversions or fusions) for which no heterotypic collision was identified in the previous 30 s were, by process of elimination, considered to constitute unaided events.

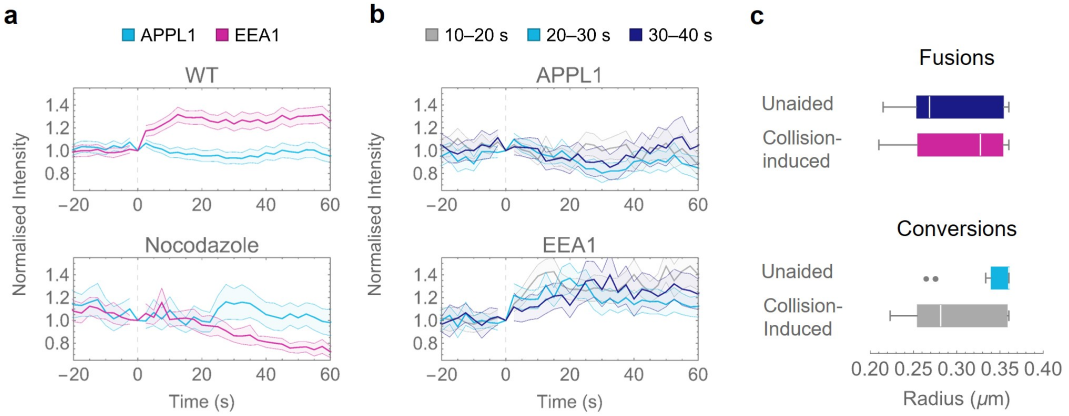


**Fig. 4. Characterization of identified events. (a)** Population-average intensity traces for wild-type (WT) (*top*) and nocodazole-treated (*bottom*) cells, aligned with the start of each conversion event. WT cells demonstrate a steady increase in EEA1 signal and decrease in APPL1 signal from the start of colocalisation, whereas nocodazole-treated cells show no clear separation between the channels, on average (i.e., detected events likely represent 'conversions' that subsequently revert). Up until the end of colocalisation, average intensity was calculated using the mask of the object detected in the APPL1 channel; afterward, it was calculated using the mask of the object detected in the EEA1 channel. Intensity as normalised by the value at the start of detected colocalisation. WT data represent $>$100 events; nocodazole-treated data represent $>$25 events; error range shows 95% confidence interval. **(b)** Average intensity traces for WT cells, separated into cohorts according to duration of conversion—short (10–20 s), medium (20–30 s), and long (30–40 s)—for APPL1 (*top*) and EEA1 (*bottom*) channels. **(c)** Size distributions of populations of APPL1 endosomes, segmented into four types of events. Endosomes that undergo unaided (direct) conversions are, on average, larger than those that undergo collision-induced conversions or fusions.

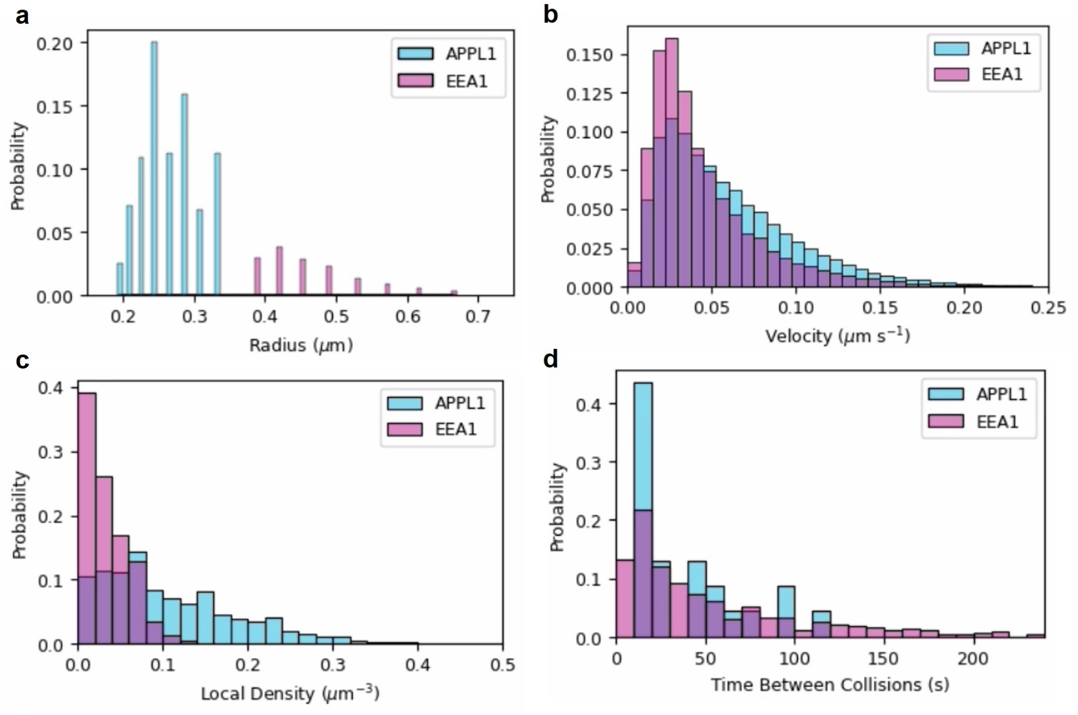


e

| Quantity ($x$) | Units | $\langle x_{APPL1} \rangle$ | $\langle x_{EEA1} \rangle$ |
|---|---|---|---|
| Instantaneous velocity | μm $s^{-1}$ | 0.15 ± 0.11 | 0.12 ± 0.09 |
| Radius | μm | 0.28 ± 0.05 | 0.42 ± 0.08 |
| Mean number density | $\mu m^{-3}$ | 0.074 ± 0.046 | 0.030 ± 0.010 |
| Mean free path | μm | 7.7 ± 3.3 | 16.1 ± 6.2 |
| Mean time between collisions; expected | s | 45 ± 15 | 148 ± 72 |
| Mean time between collisions; estimated | s | 45 ± 48 | 49 ± 48 |

f

| | APPL1–EEA1 | APPL1–EGF1 |
|---|---|---|
| Collisions (#) | 6953 | 315 |
| Collisions (# of APPL1 tracks) | 14.8% | 1.3% |
| Conversions (# of APPL1 tracks) | 2.9% | 0.0% |

**Fig. 5. Verification of collision detection.** Utilising observed distributions of endosome **(a)** sizes, **(b)** instantaneous velocities, and **(c)** local endosome densities, we calculated the expected mean time between collisions; this agrees well with **(d)** measured mean time between collisions. **(e)** Averages and standard deviations of distributions in **a–d**. The expected mean time between collisions lies well within the error bars of the estimated mean time between collisions, reinforcing that the calculated values are in line with expectations based on physical principles. **(f)** Comparison of detected collision and conversion frequencies between APPL1 and EEA1 or EGF-bearing late endosomes, respectively. Collisions between APPL1 and EGF-bearing late endosomes do occur, although at a significantly reduced frequency compared to APPL1 and EEA1 endosomes, but no 'conversions' are detected based on colocalisation of APPL1 and EGF-bearing late endosomes following such collisions.

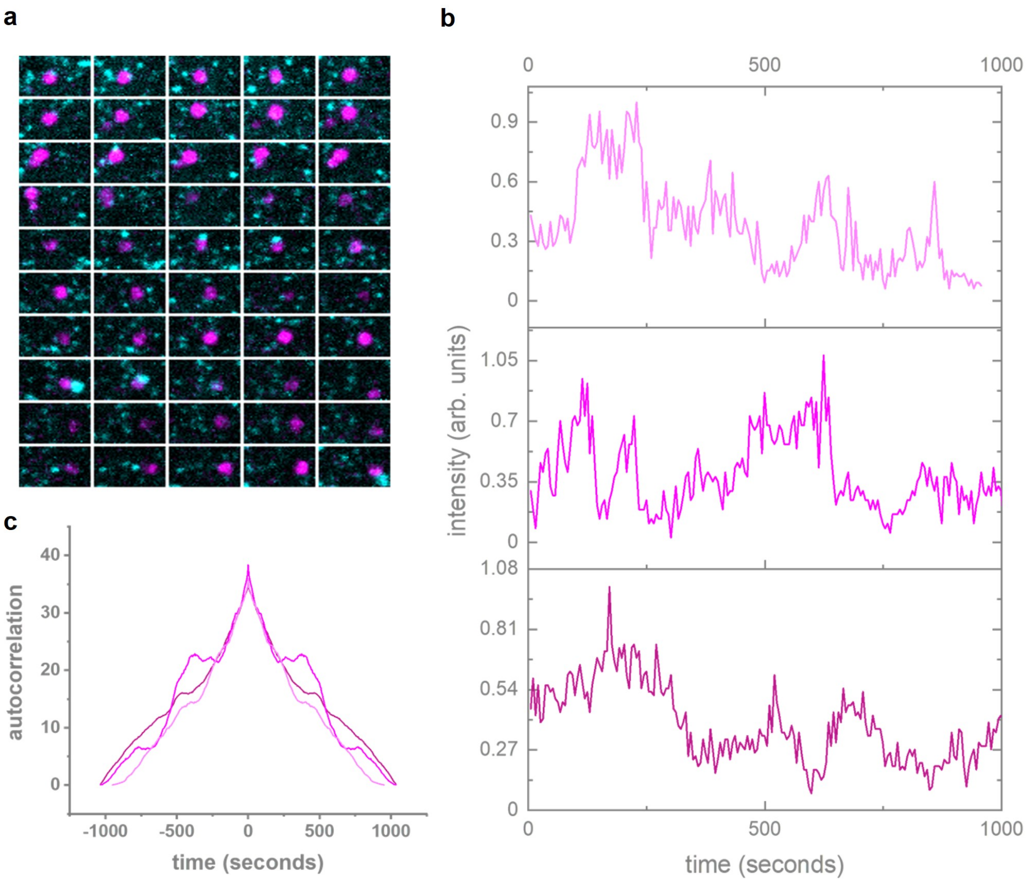


**Fig. 6. Endosomes show large EEA1 intensity fluctuations. (a)** Montage of an EEA1 positive endosomes showing fluctuating signal. **(b)** EEA1 TagRFP-T intensity traces of three representative EEA1 positive endosomes. **(c)** Autocorrelation of the intensity signals reveals secondary peaks that hint at periodicity in fluctuations of EEA1 signals.

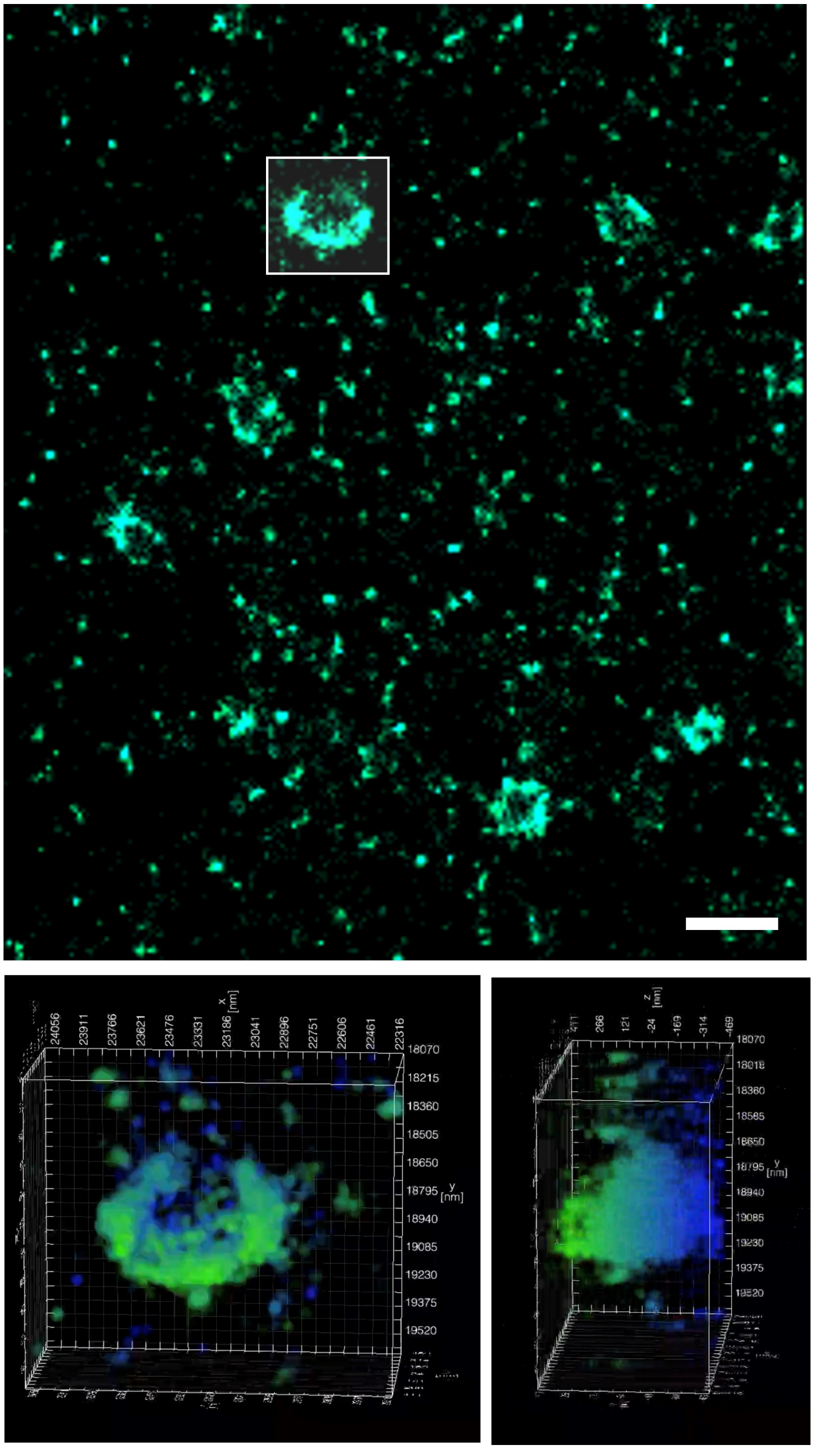


**Fig. 7. EEA1 forms distinct domains over the endosome surface.** 3D PALM imaging of Dendra2-EEA1 shown as a maximum intensity projection (*top*) and as a 3D volume (*bottom*). Intensity is colour-coded to $z$ position. Scale bar = 1 µm.

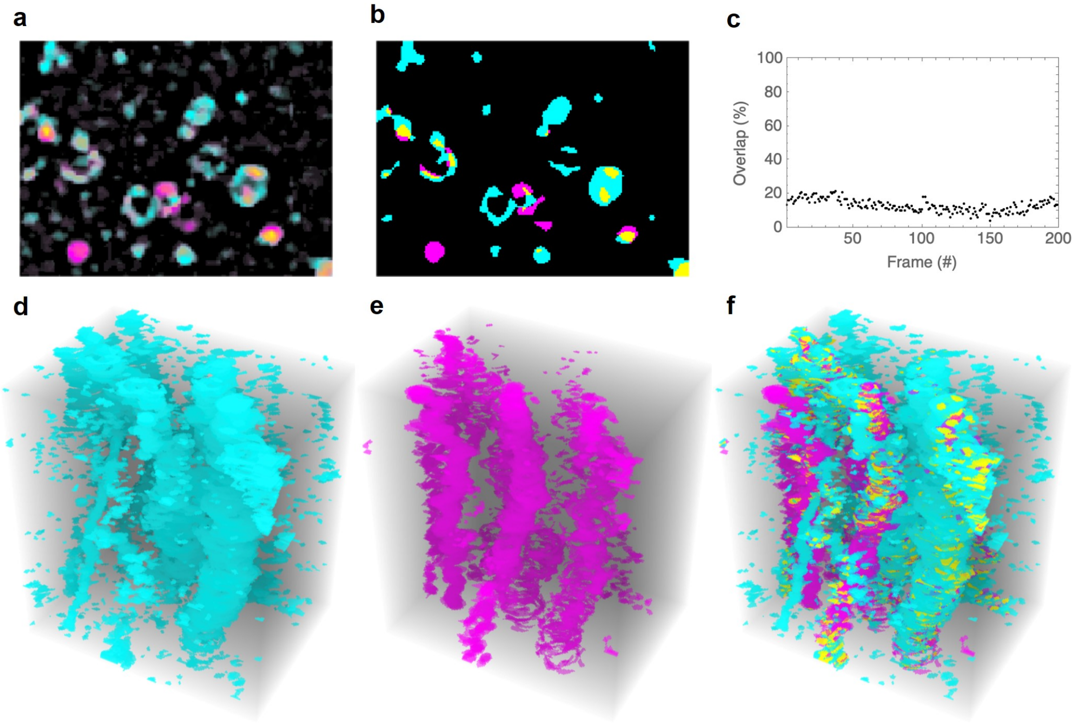


**Fig. 8. Quantification of counter-clustering.** **(a)** Sample SRRF image showing APPL1 (*cyan*), EAA1 (*magenta*), and APPL1 plus EEA1(*yellow*). **(b)** Binary mask corresponding to **a**, following thresholding to identify clusters of signal in each channel, showing pixels identified as APPL1 (*cyan*), EAA1 (*magenta*), and APPL1 plus EEA1(*yellow*). **(c)** Percentage of overlapping signal pixels per frame, calculated as total pixels containing both APPL1 and EEA1 signals divided by total pixels containing APPL1 and/or EEA1. In any given frame, the overlap of segmented APPL1+EEA1 channels, as a percentage of total signal pixels present in either the APPL1 or EEA1 channel, is $13.1 \pm 3.7\%$, indicating that approximately 87% of all clusters contain either detectable APPL1 or EEA1 signal, but not both. **(d–f)** Volume renderings of segmented images over 200 frames, showing **(d)** APPL1, **(e)** EEA1, and **(f)** APPL1+EEA1. APPL1 signal is indicated by cyan, EEA1 by magenta, and overlap (APPL1+EEA1) by yellow shading. Time is represented along the vertical axis.

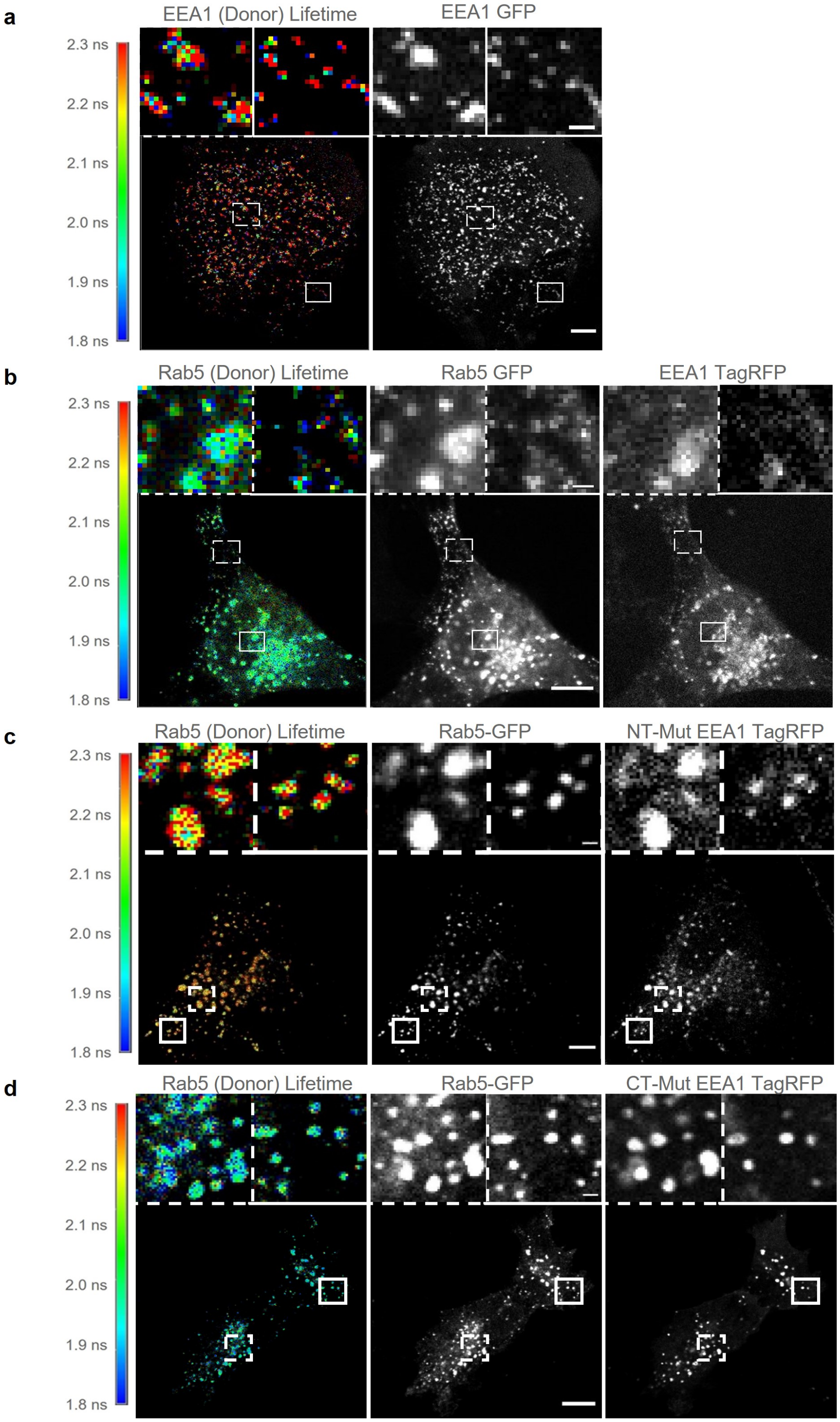


**Fig. 9. EEA1–Rab5 FRET interaction corresponds to EEA1 N-terminal binding.** Representative FLIM-FRET experiments of RPE1 cells transfected with **(a)** solely EEA1 EGFP, **(b)** Rab5 GFP + EEA1 TagRFP **(c)**, Rab5 GFP + NT-Mut EEA1 TagRFP, or **(d)** Rab5 GFP + CT-Mut EEA1 TagRFP. Coloured scale bar represents donor lifetime ranging from 1.8 ns (*blue*) to 2.3 ns (*red*). Left panels show FLIM images of donor lifetime; middle and right panels show EEA1 and Rab5 fluorescence intensity. Boxes indicate regions of zoomed inserts. Scale bar = 10 µm. Zoomed insert scale bar = 1 µm.

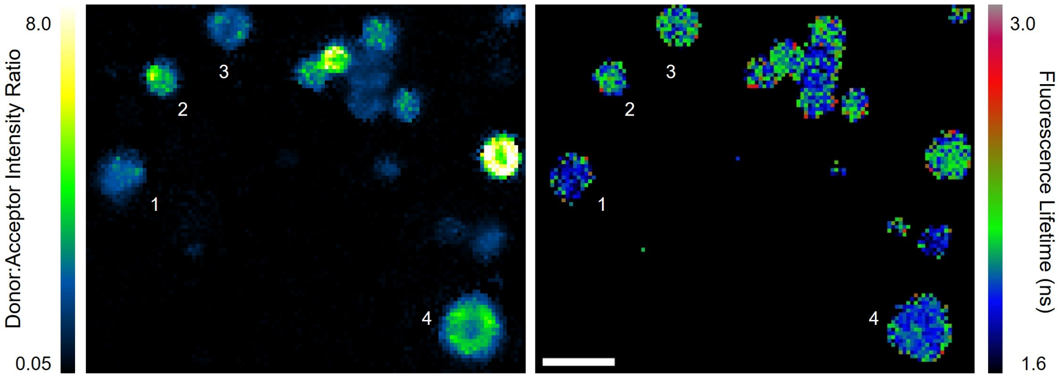


**Fig. 10. EEA1–Rab5 interaction is independent of donor:acceptor intensity ratio.** (*Left*) Representative FLIM image of RPE1 cell transfected with EEA1 EGFP and Rab5 RFP showing donor:acceptor intensity ratio. Coloured scale bar represents intensity ratio at each pixel, ranging from low D:A (*blue*) to high D:A (*white-yellow*). (*Right*) The corresponding donor lifetime is shown at each pixel, as indicated in the coloured scale bar, from 1.6 ns (*blue*) to 3.0 ns (*red*). Example endosomes are numbered: #1 has both low D:A ratio and lifetime, #2 has both high D:A ratio and lifetime, #3 has low D:A ratio but high lifetime, and #4 has high D:A ratio but low lifetime. Scale bar = 2 µm.

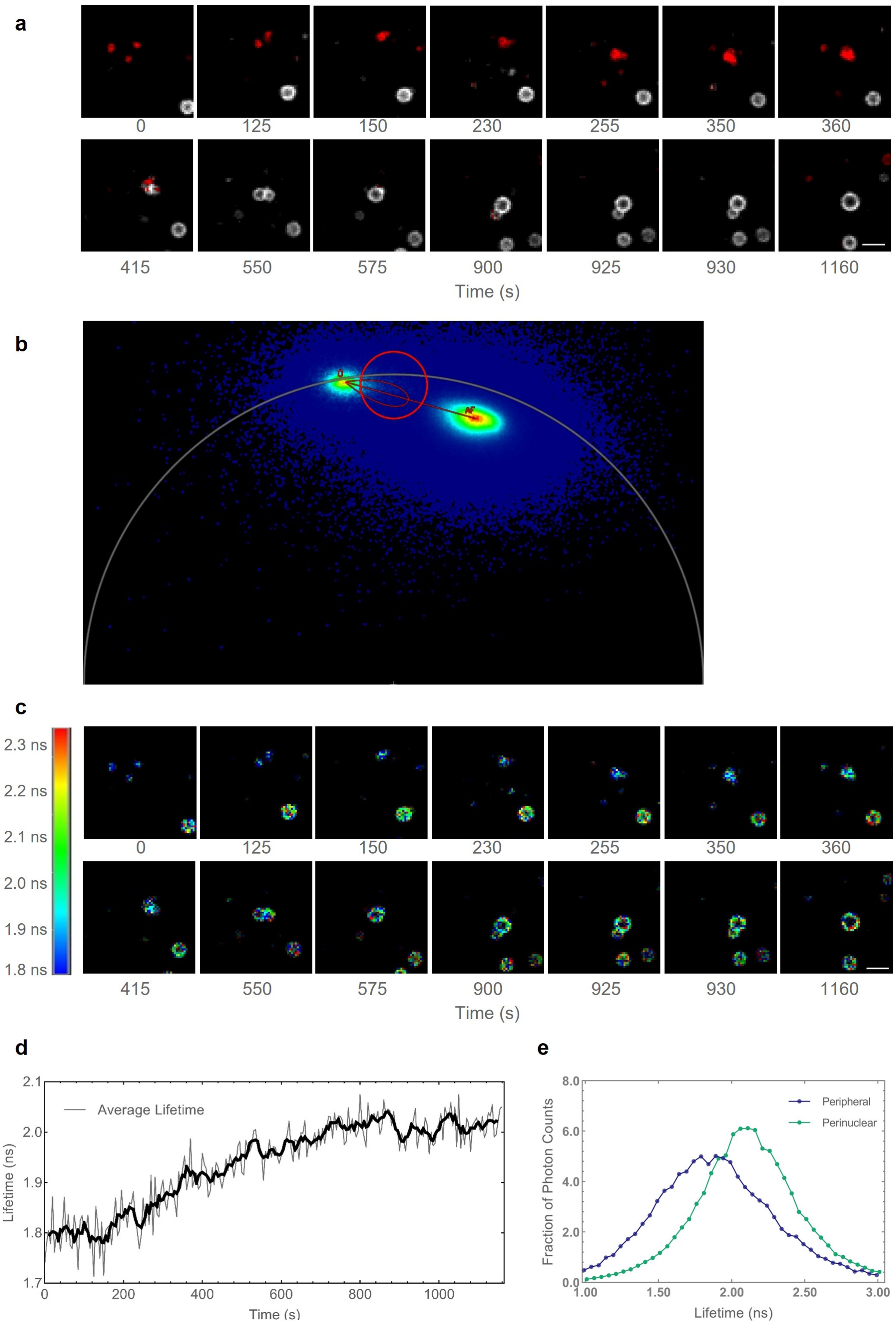


**Fig. 11. N-terminal to C-terminal EEA1 maturation captured in phasor plot and average lifetime analysis of FLIM data. (a)** Representative montage of EEA1 N- to C-terminally bound conversion (see. Fig. 3d), with pixels corresponding to FRET (*red*) and to native donor lifetime (*grey*). FRET and non-FRET pixels are assigned using the phasor plot in **b**. Scale bar = 2.5 µm; time is measured in seconds. **(b)** Phasor plot of donor (D) and acceptor (A) lifetimes. FRET trajectory is indicated by red overlay, circle corresponds to the population of FRET-associated pixels. **(c)** Representative montage of EEA1 N- to C-terminally bound conversion (see Fig. 3d). Coloured scale bar represents donor lifetime ranging from 1.8 ns (*blue*) to 2.3 ns (*red*). Scale bar = 2.5 µm; time is measured in seconds. **(d)** Average pixelwise donor lifetime of converting endosome in **c**. Lifetime is measured in ns; time is measured in seconds. Thick line corresponds to a 5-frame rolling average. **(e)** Normalised frequency histograms of the detected fluorescence lifetimes of EEA1-EGFP photons measured in live peripheral endosomes (*blue*) and perinuclear endosomes (*green*). Mean lifetimes calculated from these data are 1.91 and 2.12 ns for peripheral and perinuclear endosomes, respectively.

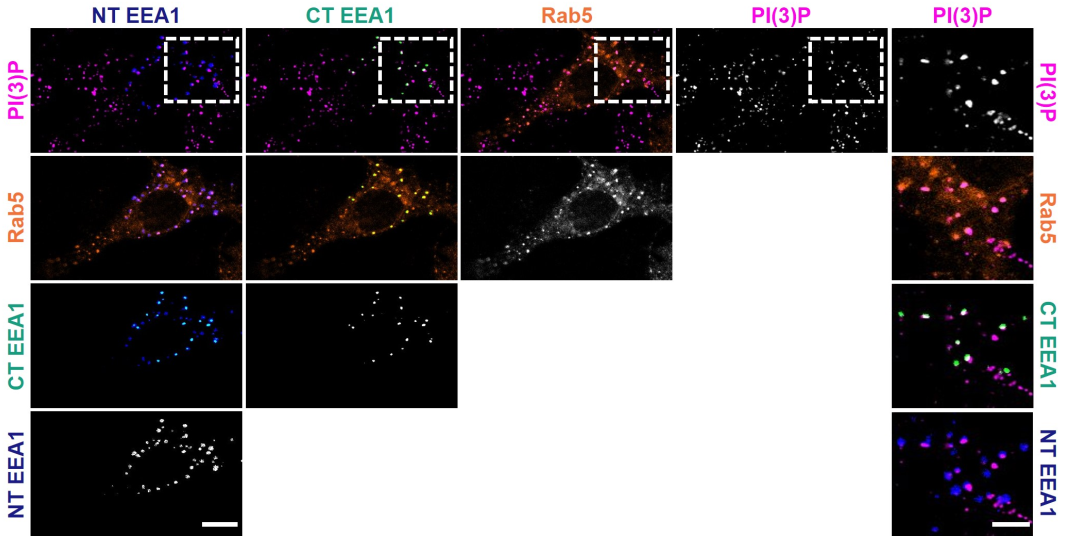


**Fig. 12. C-terminal but not N-terminal EEA1 binding colocalises with PI(3)P.** Representative image of 2xFYVE-GST PI(3)P staining with EEA1 FLIM imaging. NT and CT EEA1 channels were extracted using two-component fitting of fluorescence lifetime and each channel shown combinatorically: PI(3)P (*magenta*), Rab5 (*orange*), CT EEA1 (*green*), and NT EEA1 (*blue*). Dotted box indicates zoomed insert (rightmost column). Scale bar = 10 µm, insert scale bar = 5 µm.

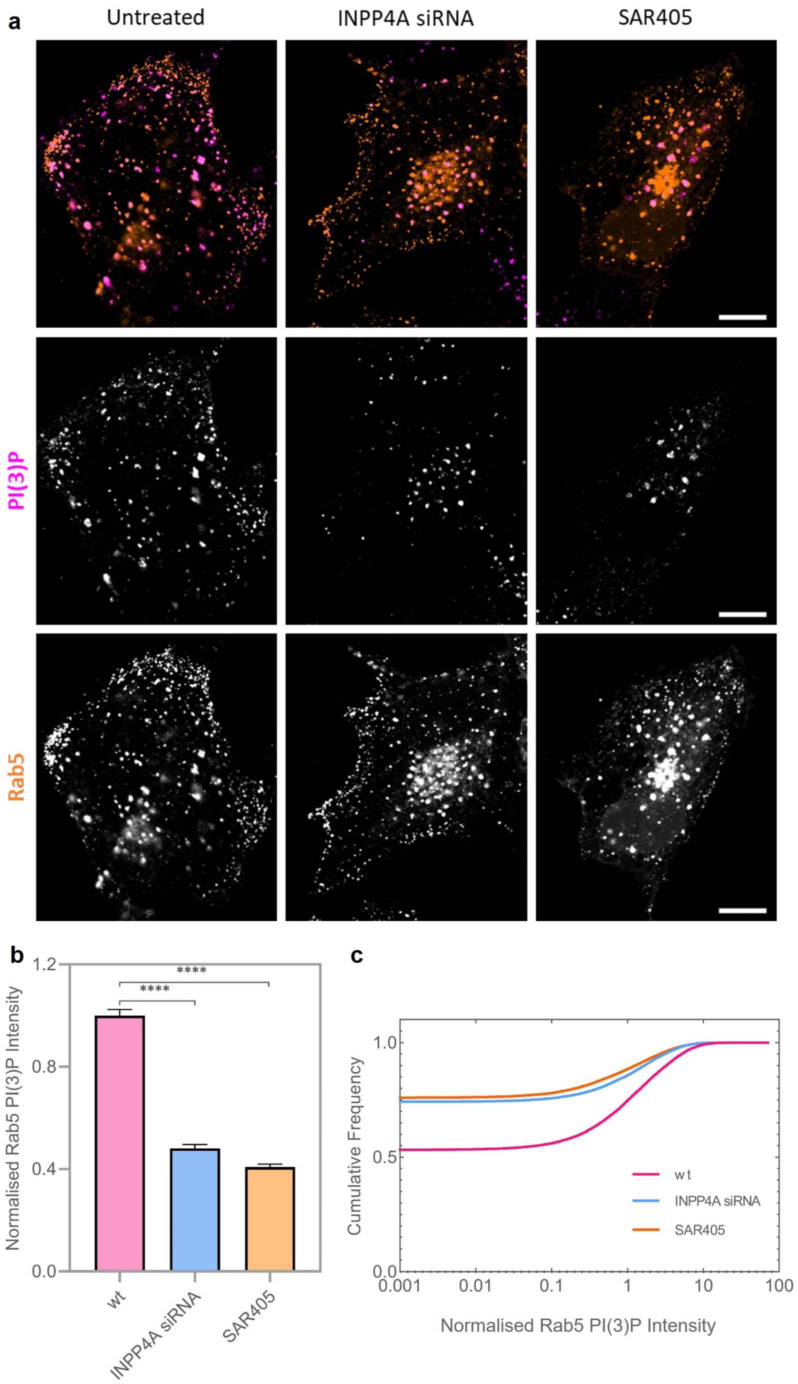


**Fig. 13. SAR405 and INPP4A siRNA lead to a reduction in PI(3)P on Rab5 endosomes.** **(a)** Representative image of Rab5 (*orange*) and 2xFYVE PI(3)P (*magenta*) staining in untreated, INPP4A siRNA, or 100 nM SAR405-treated cells. Scale bar = 10 µm. **(b)** Bar chart of normalised PI(3)P intensity present on Rab5 endosomes in untreated (*magenta*), INPP4A siRNA (*blue*), or SAR405-treated (*orange*) cells. Error bar indicates S.E.M.; $n > 50$ cells for each condition. Difference in mean values was tested using an ordinary one-way ANOVA; **** indicates $p < 0.0001$. **(c)** Cumulative frequency histogram of **b**, showing untreated (*magenta*), INPP4A siRNA (*blue*), SAR405-treated (*orange*) cells.

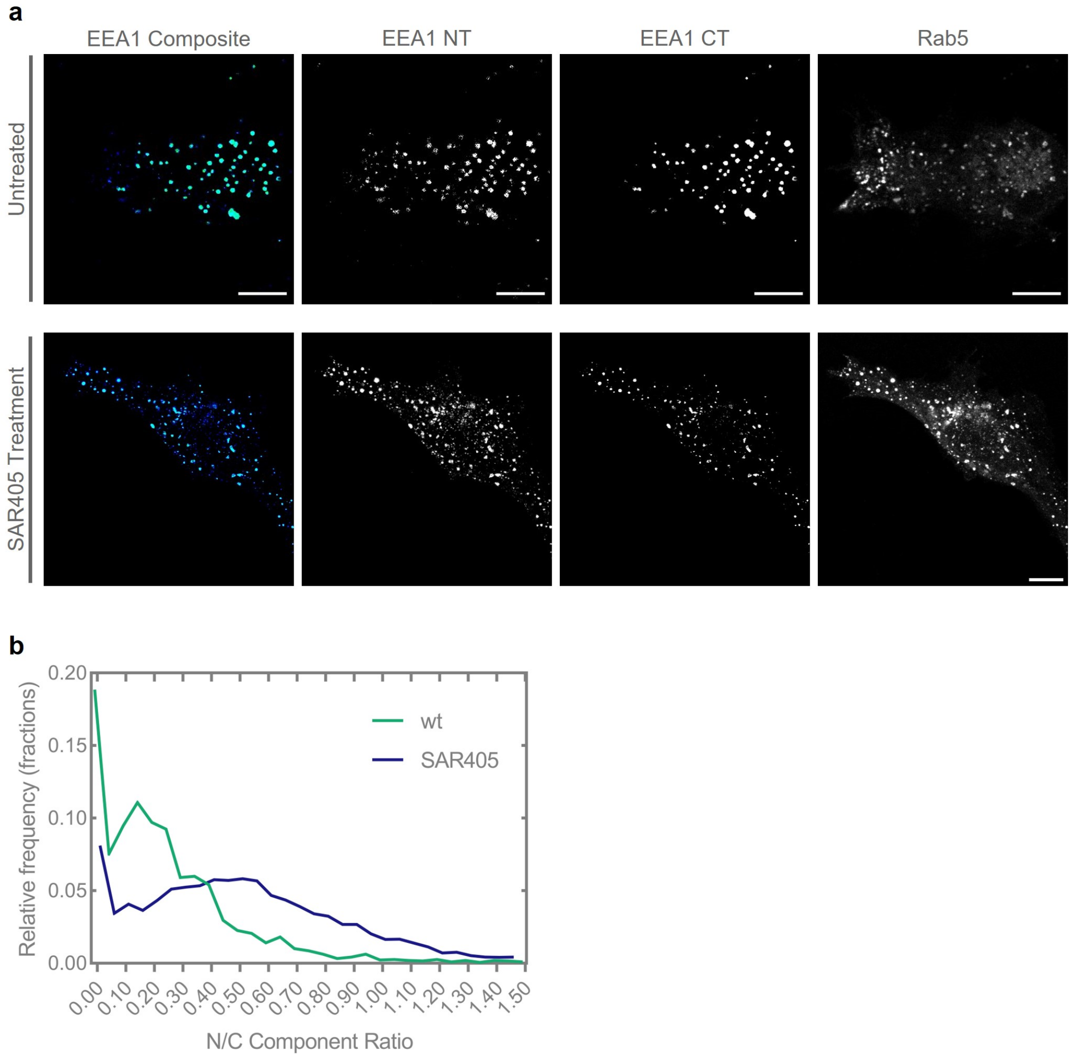


**Fig. 14. SAR405-treated cells show reduced but detectable CT EEA1 binding. (a)** Representative image of EEA1-EGFP Rab5-RFP expressing cells, either untreated (*top*) or SAR405-treated (*bottom*). NT EEA1 (*blue*) and CT EEA1 (*green*) were separated using two-component fitting. Scale bar = 10 µm. **(b)** Histogram of endosomal pixels in **a** with wild-type (*green*) and SAR405-treated (*blue*) cells; showing the ratio of N- to C-terminal lifetime components. The N- and C-terminal components were fit as described in Fig. 3 with $\tau_1$ = 1.006, $\tau_2$ = 2.600 ns.

### Supplementary Note 1: Supplementary Notes

#### Supplementary Note 1.

***Simulating a single endosome.*** The grid that represents the endosome's surface forms a Fibonacci Sphere in which neighbouring grid points are approximately equidistant. Each grid point has two layers, which can be empty or occupied by nodes. The outer layer can be occupied by either PI(3,4)P2 or PI(3)P, while the inner layer can be occupied only by Rab5. Movement of a node over the endosome's surface consists of moving from its original grid point to a neighbouring empty grid point on the same layer. Each node can be either free or occupied, with one of the following three agents attached to it: (i) APPL1, which can attach to a grid point that has unoccupied PI(3,4)P2 and Rab5 nodes, and occupies both nodes on that grid point; (ii) INPP4A, which can attach to and occupy PI(3,4)P2; and (iii) EEA1, whose N-terminus can attach to and occupy Rab5, whilst the C-terminus can only attach to a grid point that has unoccupied PI(3)P and Rab5 nodes, and occupies both nodes on that grid point.

Nodes have a certain probability (rate) of moving to an empty grid point in the neighbourhood. This rate is smaller for heavier nodes and decreases if there is an agent attached to the node (modelling the fact that diffusion speed is slower for a larger mass). We have modelled clustering of nodes in the following manner: for any type of node, (i) the rate of moving decreases with increasing number of nodes of the same type in the neighbourhood, and (ii) the rate of moving to a particular empty grid point in the neighbourhood increases as the number of nodes of the same type increase in the neighbourhood of that grid point. Agents have specific probabilities (rates) of attaching to or detaching from specific nodes. Agent clustering was implemented by increasing attachment rates for an agent to a node according to the number of agents of the same type in the neighbourhood of that node.

It was experimentally observed that levels of EEA1 stochastically rise and drop sharply throughout the time course, which is reminiscent of mass attachments and detachments. To model this phenomenon, we have enabled EEA1 to detach in clusters in our simulation. This was implemented by making the rate of detachment of EEA1 from a node increase with the number of EEA1 detachment events that happened in the recent past in the neighbourhood of that node. When INPP4A is attached to a PI(3,4)P2 node, it has a certain probability (rate) of converting that PI(3,4)P2 to PI(3)P. Immediately after conversion, INPP4A detaches from the PI(3)P. At this instant, there is a certain probability that this INPP4A will jump to a free PI(3,4)P2 node that lies within a certain distance (labelled jump distance) of the node it was originally attached to.

For all the simulation runs, we start out with a predetermined number of nodes of each type, which are clustered together according to their types. These clusters are arranged randomly on the surface of the endosome. We start out with no agent attached to any of the nodes. The conversion time of the endosome is measured as the first passage time of the fraction of PI(3,4)P2 converted to PI(3)P crossing a fixed threshold.

***Simulating collision of two endosomes.*** The collision of two endosomes (one young and one mature) occurs over a short finite interval of time. In this time interval, there is a pool of a given number of EEA1 from the mature endosome surrounding the young endosome that attach whenever any free node of the appropriate type is available. Whenever any EEA1 stochastically detaches from the young endosome, it goes to the EEA1 pool surrounding it instead, and is added back whenever any free node is available. If there are an excess of free nodes, the free nodes to which EEA1 will attach are randomly selected with equal probability. N-terminus attachments are given precedence over C-terminus attachments because we assume the collision is occurring with a mature endosome; on mature endosomes, most EEA1 attachments are observed to occur via the C-terminus, leaving the N-terminus free.

***Simulating bulk dynamics.*** We first simulated the effect of one or multiple collisions at various times since initiation on the conversion time distribution. Then, we simulated a system in which new endosomes are introduced at fixed intervals of time, whilst old ones exit at the same intervals (to keep the population of endosomes in the system constant). The endosomes in the system are free to collide randomly. The conversion time of endosomes which did not experience any collisions is drawn from the previously obtained distribution of conversion times of endosomes not undergoing collision. Similarly, endosomes which underwent a given number of collisions at given times (where only collisions with already converted endosomes are counted) have their conversion times drawn from the appropriate previously obtained distributions. Endosomes for which the conversion time was longer than the duration of their presence in the system were labelled as unconverted.

***Simulating fusion of two endosomes.*** We have simulated fusion of two endosomes of different ages by fusing the bottom pole of the first endosome with the top pole of the second one. Thus, nodes near the top of the first endosome stay clustered near the top even after fusion but the nodes near the bottom pole are spread out just above the equator. The second endosome displays the inverse behaviour after fusion, with nodes near the bottom remaining clustered near the bottom, but those near the top spread out just below the equator. The longitudes of the nodes are roughly maintained before and after the fusion. This ensures that the nodes which were clustered before the fusion still remain roughly clustered after the fusion. The total size of the endosome increases after fusion such that the total surface area is conserved (i.e., the surface area of the fused endosome is equal to the sum of surface areas of the first and second endosomes). This ensures that the distance between the neighbouring nodes is roughly the same before and after the fusion.
